# Metabolic modeling and functional genomics reveal taxa and host gene interactions in colorectal cancer

**DOI:** 10.64898/2026.01.26.700635

**Authors:** Katja Della Libera, Elizabeth M. Adamowicz, Amanda Muehlbauer, Sambhawa Priya, Adnan Alazizi, Francesca Luca, Ran Blekhman

**Author notes:** These authors contributed equally to this work (Katja Della Libera, Elizabeth M. Adamowicz).

## Abstract

Colorectal cancer (CRC) is associated with changes in the microbial communities in the tumor microenvironment. Although metabolic reprogramming is an important feature of host cells in CRC, little is known about metabolic changes in the tumor-associated microbiota and how these microbial metabolic alterations can contribute to disease. Here, we investigated metabolic host-microbiome interactions in CRC using complementary computational and experimental approaches. Using patient-specific *in silico* metabolic models across three independent datasets, we discovered that *Fusobacterium*, a cancer-promoting taxon, consistently grows faster in tumor-associated versus normal tissue-associated microbiomes. This finding prompted us to investigate whether host metabolic changes drive these microbial growth advantages. By integrating our metabolic predictions with host transcriptomics data, we identified correlations between tumor gene expression and the growth of CRC-associated taxa (including *Porphyromonadaceae*, *Blautia*, and *Streptococcus*), as well as associations between host genes and microbial metabolism of dietary components (including choline, amino acids, and starch). To test whether these correlations reflect causal relationships, we simulated spent medium experiments *in silico*, demonstrating that *Blautia* preferentially grows on metabolites produced by tumor versus normal host cells. We further validated the direct impact of microbes on host metabolism using an *in vitro* system, where colon cancer cells exposed to human microbiomes showed gene expression changes in response to specific taxa including *Bilophila*, *Anaerotruncus*, and *Escherichia*. Together, these findings reveal a metabolic dialogue between host and microbiome in CRC, where tumor metabolic reprogramming creates a favorable environment for pathogenic microbes, which in turn may reinforce tumorigenic processes through metabolic crosstalk.

## Introduction

Colorectal cancer (CRC) is the third most common cancer and the second most common cause of cancer-related deaths worldwide [1]. The human colon is estimated to host approximately 10^14^ bacterial cells, or 70% of the total gut microbiome bacterial load [2]. Numerous studies link perturbations in the composition and function of the colonic microbiome to the development and progression of colorectal cancer; however, the specific causal role of the microbiome as a whole community, as well as the individual roles of its constituent species, remains a debated topic in the field. Deciphering these causal effects will require a detailed understanding of how the various taxa that make up the gut microbial community change in abundance and metabolic activity within the tumor microenvironment.

Metabolic reprogramming is a well-established feature of CRC carcinogenesis, as rearrangement of cellular metabolism is required to sustain the high rates of cell division and angiogenesis required for CRC [3]. Changes in human host cell metabolism include an increase in glycolysis activity and pentose phosphate pathway activity; increased lactate oxidation; increased nucleotide and lipid metabolism; and decreased oxidative phosphorylation and serine biosynthesis [4]. The Warburg effect, wherein cancer cells preferentially ferment their glucose supply into lactate rather than shuttling it into mitochondrial oxidative phosphorylation, is a feature of colorectal cancer development and may be induced by hypoxic conditions within a tumor microenvironment or mutations in oncogenic signaling pathways [5–7]. These and other metabolic changes in diseased cells can have complex reciprocal interactions with associated microbial communities, as metabolic products from microorganisms may serve as metabolic precursors or gene regulatory signals for host cells (and vice versa). For example, butyrate and other short chain fatty acids produced by microbial fermentation are the main carbon source for colonocytes, and hundreds of genes in the human GI tract are differentially regulated by the microbiome [8].

CRC is associated with a variety of microbiome community-level changes, such as a decrease in overall diversity [2,9,10] and an increase in hydrogen sulfide-producing organisms [11]. Three taxa in particular have been causally linked to colorectal carcinogenesis: *Fusobacterium nucleatum*, genotoxic *Escherichia coli*, and *Bacteroides fragilis* [12]. Numerous studies have reported elevated abundances of these species in CRC compared to healthy tissues, and many of their metabolic functions have been implicated in disease onset and outcome. A variety of mechanisms can mediate these effects; metabolites produced by these organisms may induce mutations [13], promote inflammation [14–18], upregulate oncogene pathways [19–22], and enhance chemoresistance against common therapeutics [23]. Other taxa are protective against CRC development and progression. *Clostridium butyricum*, for example, has been reported to inhibit tumor development in genetically susceptible mice, and to alter the composition of the gut microbiome, decreasing the abundance of pathogenic bacteria such as *Desulfovibrio* while increasing the abundance of beneficial microorganisms such as the short chain fatty acid producing Ruminococcaceae. However, disentangling the complex ecological relationships between protective and potentially carcinogenic microbiome taxa and the host remains an ongoing challenge in the field.

Microbiome effects on the expression of host genes influencing host metabolism and immunity may be particularly relevant for CRC. Differential expression of host microRNAs (miRNAs) in CRC is associated with microbiome composition [24]. Pathogenic bacteria such as *Helicobacter pylori*, *Salmonella enterica* var. *Typhimurium,* and *Listeria monocytogenes* can also activate host mRNAs, which reduce the host immune response and increase autophagy [8]. One study examining host gene expression and microbiome composition across several gastrointestinal diseases identified a common set of host inflammatory and energy metabolism genes correlated with disease-specific microbes [25]. While many associations between microbiome composition and host gene expression have been identified, the causal direction of these relationships remains unclear [8]. To study these causal relationships, an *in vitro* approach has been developed that characterizes gene expression responses to microbial communities in host colonic cell cultures [26,27]. This *in vitro* approach has revealed variation in the expression of over 6,000 genes at various time points after co-culture with a gut microbial community, including enrichment of genes associated with conditions such as obesity and colorectal cancer [26]. Many studies exploring the host gene expression response to the microbiome focus on microbial composition but do not address metabolic activity; this is mainly due to the difficulty of studying metabolism in the colonic microbiome environment, where approaches such as untargeted metabolomics are unable to distinguish between host-derived, diet-derived, and microbiome-derived metabolites, as well as how changes in one side might impact the other.

One way to circumvent the challenges of addressing causal questions around microbial metabolism in CRC is through mathematical modeling. Models based on experimentally determined microbial growth patterns can predict how particular taxa within a sample microbiome will use and generate metabolites from a medium. This approach sidesteps the challenge of disentangling the sources of metabolites identified by *in vitro* or *in vivo* sampling, enabling researchers to generate hypotheses about the relationship between microbial metabolism and human gene expression that can be tested experimentally. Flux balance analysis (FBA) is a particularly useful mathematical approach for modeling activity within biochemical networks [28]. Genome-scale metabolic models can quantify the metabolic reactions performed by an organism (as identified by enzyme-encoding genome sequences) and use FBA to predict the metabolites produced and consumed by the organism’s biochemical network when an objective function, usually biomass, is maximized [28]. Prior studies have used FBA-based models to predict the metabolic activity of all microbes within a community based on a defined metabolic growth environment (such as that of the colon) [29]. In addition, by modifying the growth environment, dietary interventions or changes in surrounding host cell metabolism can be considered.

The growing use of metabolic modeling approaches in cancer research has focused primarily on cancer cells despite the well-established role of microbial metabolism in CRC. A recent preprint demonstrated the value of understanding metabolic changes in the microbial community as a whole in colorectal cancer by showing differential response in a simulated diet experiment [30]. The goal of our study is to use *in silico* models of microbiome communities to identify specific metabolic features of the microbiome likely to play significant functional roles in CRC. We built these communities for three colorectal cancer microbiome datasets from different patient sample populations; all three datasets were generated using 16s rRNA sequencing of tumor and adjacent normal tissue colonic biopsy samples from CRC patients [31–33]. We used the microbiome community metabolic modeling package MICOM [34] to build site- and patient-specific *in silico* microbiomes, and used these communities to examine predicted microbial growth rates and metabolic flux across a shared growth medium/diet environment. For one of the datasets [31], we also examined correlations between host gene expression and microbial metabolic activity using LASSO penalized regression and built gene-expression-based *in silico* metabolic models for the human host cells using CORDA (Cost Optimization Reaction Dependency Assessment) [35]. By using these host cell models for spent media experiments we further tested whether the host cell metabolism in cancer or normal tissue influences the metabolic activity of the microbiome and vice versa.

Finally, we complement our *in silico* approach with an *in vitro* study of gene expression in colonic epithelial cells treated with different gut microbiomes, identifying genetic responses with RNAseq and metagenomic shotgun sequencing [26,27]. This approach has not yet been used on different cell lines to compare the impact of the microbiome on tumor vs normal tissue. Here we characterize the response of epithelial cells to the abundance of specific microbial taxa *in vitro* and compare it to significant correlations we found in our paired dataset in [31]. This work provides new insights into how microbial metabolism changes in CRC compared to normal tissue-associated microbiomes, and how changes in host gene expression are associated with microbiome metabolism.

## Results

### In silico microbiome communities predict differences in growth rate and metabolic activity of taxa in tumor compared to normal tissue-associated communities

We used 16s rRNA microbiome taxonomic abundance data from paired tumor-normal patient biopsy samples across three previously published studies [31–33] to build *in silico* microbiome communities specific to a given patient and site (tumor or normal). We will hereafter refer to the datasets generated from these three studies as Burns2015, Hale2018, and Niccolai2020, respectively. We then used these *in silico* communities to perform flux-balance analysis, which predicts the growth rate of a given taxon in a community, as well as the activity (flux) of all metabolic reactions that taxon will perform based on a given medium environment (see **Methods**). Within each study population, we identified taxa with significantly different predicted growth rates between tumor and normal tissue-associated microbiomes, as well as metabolic exchange pathways (reactions which allow metabolites to enter and exit the modeled cell) with significantly different predicted fluxes between tumor and normal tissue associated microbiomes. We then compared these taxa and exchange flux pathways across study populations to identify common signatures in CRC. An overview of our study approach can be found in **Figure 1A**.

**Figure 1:**
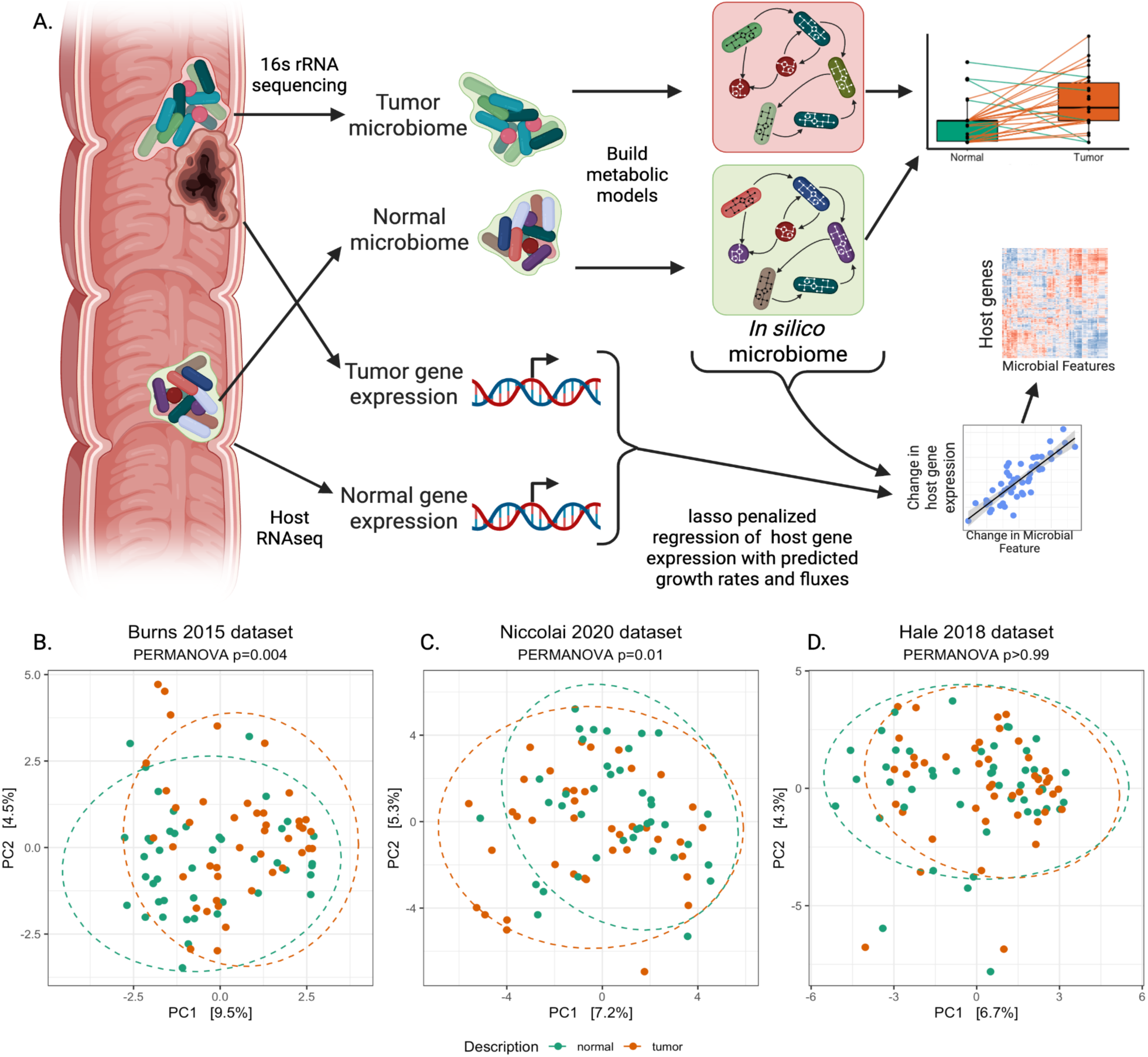
Experimental overview. **A.** Using three unique datasets of 16s rRNA microbiome data obtained from colonic biopsies of tumors and normal adjacent tissue, we built *in silico* metabolic models of each patient- and site-specific microbiome. One dataset (Burns et al. 2015) also contained host RNA-seq data from tumor and adjacent normal sites, which we used to build metabolic models of host cells as well, and correlated with microbial flux. **B-D.** Microbiome composition beta diversity (Aichison’s distance) of communities used to build *in silico* metabolic models for Burns 2015 (**B**.), Hale 2018 (**C.**) and Niccolai 2020 (**D.**) datasets. Each dot represents a sample with tumor and normal samples represented by red and green dots, respectively.

Prior to building microbial community models, we filtered the taxonomic data from each study to remove potential contaminants and taxa below a minimum abundance threshold of 0.1% (see **Methods**), then examined differences in microbiome composition between tumor and normal sites for the resulting taxonomic tables. Alpha diversity (as measured by Chao1, Shannon, and Simpson indices) did not differ significantly between tumor and normal samples in any dataset (**Supplementary Figure 2,** lowest p-value: 0.051, Chao1 for Niccolai2020). We observed varying differences in the beta diversity between tumor and normal samples (as measured by Aitchison’s distance) for each of our datasets. While the Burns2015 dataset (**Figure 1B**, PERMANOVA *p*=0.004,) and the Niccolai2020 dataset (**Figure 1C**, PERMANOVA *p*=0.01) show significant differences in beta diversity between tumor and normal samples, the variation explained is relatively small (7-9%). The Hale2018 dataset did not show a significant separation of tumor tissue-associated and normal tissue-associated communities (**Figure 1D**, PERMANOVA *p*>0.99).

The process used to build *in silico* microbial communities is outlined in **Figure 2A** and further details can be found in the **Methods**. Briefly, we first matched each operational taxonomic unit (OTU) to a previously published metabolic model from one of two available databases [36,37]. In cases where multiple models matched a given OTU, we kept all models and merged them using MICOM [34], a community-level metabolic modeling program that we also used for subsequent analyses. We then used MICOM to combine these models into a microbial community model scaled by the relative abundance of each OTU in the model-matched OTU table (see **Methods**).

**Figure 2:**
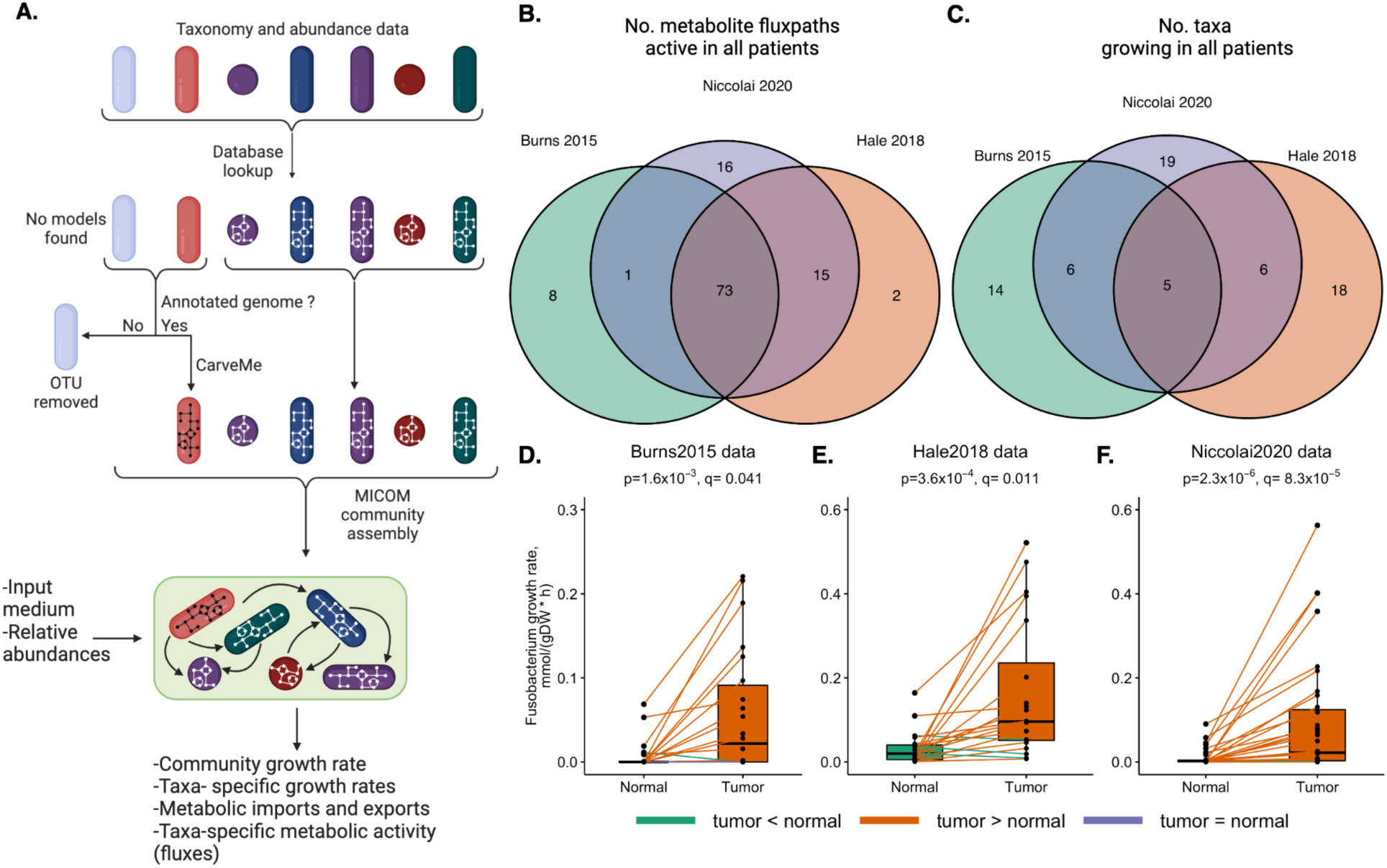
Growth rates of *Fusobacterium nucleatum* as predicted by flux-balance analysis in three different datasets. **A.** Schematic of building *in silico* microbial communities (see Methods). Briefly, we used MICOM to create site- and patient- specific *in silico* microbiome communities from previously published 16s rRNA sequencing datasets. We then performed flux-balance analysis to investigate the differences between modeled microbial growth and metabolic activity in tumor vs normal tissue associated samples. **B.** Counts of bacterial metabolite exchange fluxes (fluxpaths) active in all samples in each of the three datasets examined. **C.** Counts of bacterial taxa growing in at least half of the samples in each of the three datasets. **D-F.** Growth rate of *F. nucleatum* across all three datasets (Burns, Hale, and Niccolai, respectively). P and q values from paired Wilcoxon signed rank tests. *F. nucleatum* was the only taxon with significantly different growth in tumor vs. normal samples; no metabolite fluxpath had significantly different growth in tumor vs. normal samples across all three datasets.

Once the community models were built, we defined a shared growth medium for all communities based on a previously published set of dietary components that make up a European Union standard diet as an approximation for a western diet (see Methods) [38]. We then ran flux-balance analysis of these community models in MICOM with the standardized diet, and examined differences in community growth rate, individual species’ growth rates, and exchange flux values for tumor vs normal tissue associated communities. We found that predicted community growth rates differed significantly between tumor and normal samples in the Burns2015 dataset, wherein tumor-associated community growth was predicted to be lower than that of normal tissue-associated community growth (**Supplementary figure 3A**, paired Wilcoxon signed-rank test *p*=0.0407). For the other datasets, we did not predict a significant difference in community growth rates (Hale2018 and Niccolai2020 data in **Supplementary figure 3B** and **C**, respectively; paired Wilcoxon signed-rank test *p*=0.0990 and *p=* 0.0625 respectively).

We examined the metabolite exchange flux pathways which were differentially active between normal and tumor tissue communities. In our models, exchange flux pathways represent import and export reactions for the cell, and can therefore be thought of as a representation of the uptake and export of metabolites between the cell interior and the exterior metabolic environment. Although 73 exchange pathways differ in flux between tumor and normal samples in all three studies (see **Figure 2B**), none of these differences were statistically significant after FDR correction of Wilcoxon signed-rank tests (*q*-value<0.05). **Supplementary table 8** shows the full table of results for this analysis.

Next, we examined the taxa which were predicted to have different growth rates in tumor-associated communities as compared to normal tissue-associated communities. Of the five taxa shared between all three studies (shown in **Figure 2C**), only *Fusobacterium* was predicted to have a statistically significant difference in growth rate across communities derived from all three studies (**Figure 2D-F**, paired Wilcoxon signed-rank test with Benjamini-Hochberg FDR correction applied; **Figure 2D**, Burns2015 dataset *q*-value = 0.041; **Figure 2E**, Hale2018 dataset, *q*-value = 0.011; **Figure 2F**, Niccolai2020 *q*-value = 8.3 x 10^-5^). The full table of results can be found in **Supplementary table 8**.

Since *Fusobacterium* was predicted as the only taxon with a significantly different growth rate between tumor and normal communities in all of our datasets (**Figure 2D-F**), we further examined the abundance and *Fusobacterium*-specific metabolic activity to find pathways of interest. We found that, across all three datasets, taxa in the *Fusobacterium* phylum had a significantly higher log_2_fold change in abundance between normal and tumor associated microbiome communities (**Supplementary figure 5A-C,** paired Wilcoxon signed-rank test test with Benjamini-Hochberg FDR correction; **Supplementary figure 5A**, Burns2015 dataset *q*-value = 0.0463; **Supplementary figure 5B**, Hale2018 dataset, *q-*values 2.31x10^-9^ to 9.39x10^-4^; **Supplementary figure 5C**, Niccolai2020 dataset, *q-*values 4.54x10^-8^ to 0.031). We also found that predicted valine import was higher in *Fusobacteria* present in tumor as compared to healthy tissue in all three datasets (**Supplementary figure 5D-F**). While the differences in valine import for the Burns2015 dataset (**Supplementary Figure 5D**, paired Wilcoxon *q-*value = 0.0626), the Hale2018 dataset (**Supplementary Figure 5E**, paired Wilcoxon *q-*value = 0.301), and the Niccolai2020 dataset (**Supplementary Figure 5F,** paired Wilcoxon *q*-value= 0.0697) were not significant, all three show the same trend of increased exchange in the cancer microbiome.

### Change in host gene expression correlates with change in growth rates and metabolic activity of *in silico* microbiome communities

We next investigated whether predicted changes in microbial metabolic activity correlated with host gene expression patterns, potentially revealing mechanistic links to CRC pathogenesis. For this purpose, we used paired microbiome and host gene expression data available for one of our datasets (Burns2015) [25], enabling us to systematically identify microbe-host gene associations using lasso penalized regression [39]. We built regression models for a list of 2137 host genes pre-filtered for relevance to metabolic interactions based on the HumanCyc database (see Methods) [40]. After filtering the microbial features for prevalence (see Methods) we considered 138 metabolic fluxes with predicted differences between normal and tumor tissue across the microbial community as well as 27 taxa with predicted differences in growth rate between tumor and normal tissue for our LASSO regression model.

Using this approach we identified 43 associations between predicted taxon growth rates and host gene expression, representing five taxa associated with 43 host genes (**Figure 3A**; LASSO regression, q-value < 0.05). Notably, increased growth of Porphyromonadaceae, a family with known association to increases in CRC risk (particularly from *Porphyromonas gingivalis*) [41], was associated with upregulation of 23 host genes. These included the inflammation regulating *MTMR3*, *DNMT3A*, *PARP14*, *KDM5B*, *DGKA*, and *MAN2A1*, and the known tumor-promoting *CBR1*. Other taxa whose growth showed associations with host gene expression changes were the genera *Streptococcus* and *Blautia*, and the family Clostridiaceae, all of which are known colorectal-cancer associated taxa [10].

**Figure 3:**
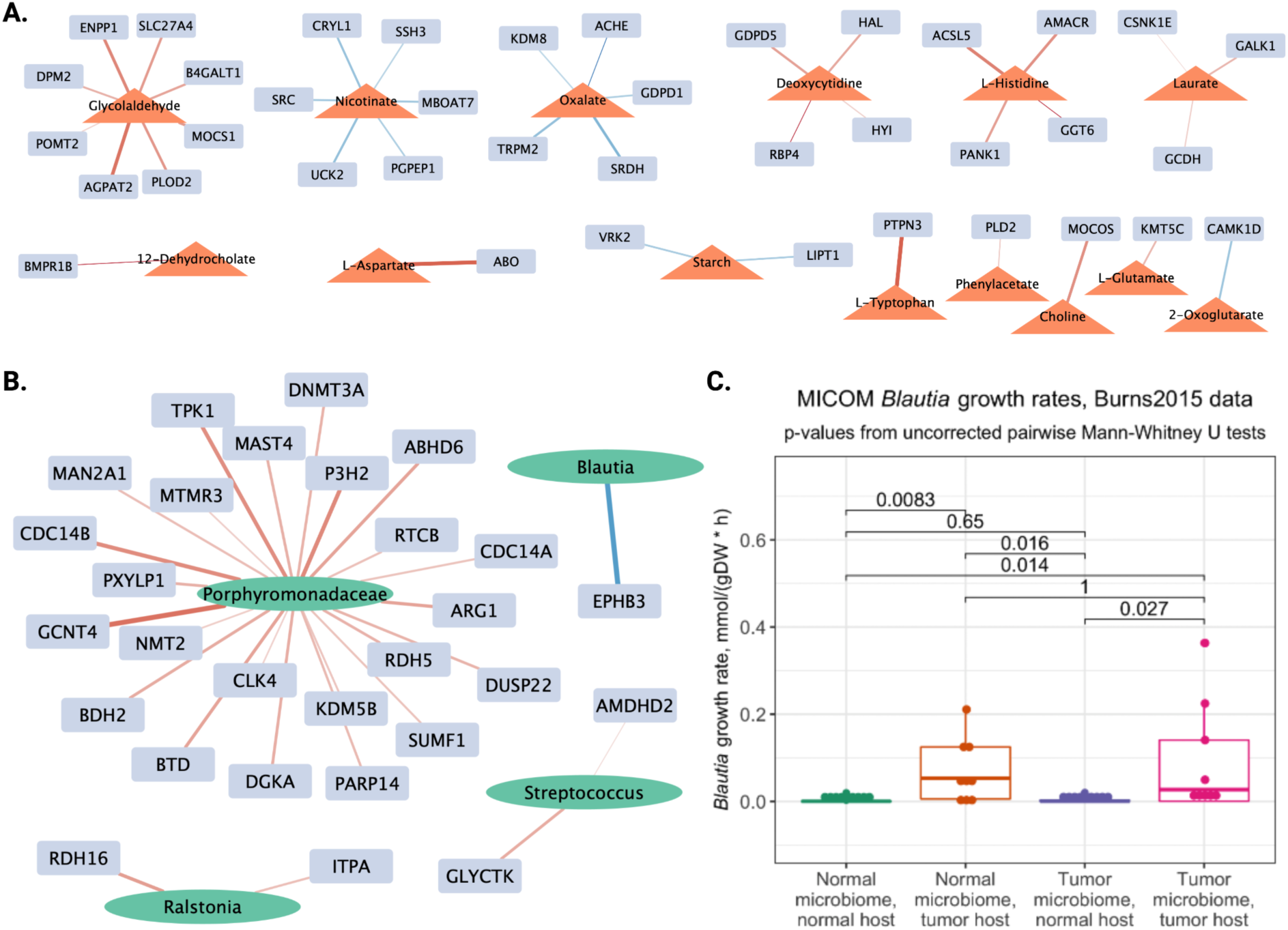
Microbiome taxa and metabolic fluxes correlating with gene expression. Networks of LASSO regression results showing **A.** significantly correlated (p-value<0.05) microbial metabolic exchanges (orange triangles) with differentially expressed genes (rectangles) and **B.** significantly correlated (p-value<0.05) microbial growth rate (green ovals) and differentially expressed genes (grey rectangles). Red lines show positive correlations, blue lines negative and the thickness of the lines indicate the size of the correlation coefficient. **C.** Further investigating *Blautia*, this taxon also shows an increase in growth rate when grown on spent-media from tumor cell models compared to spent media from normal host cell models.

In addition, our analysis identified associations between 39 host genes and 14 metabolite fluxes (**Figure 3B**). The metabolites whose uptake or production in the microbiome we found to correlate with regulation of host metabolic genes include typical signatures of the needs of tumor metabolism, such as a number of amino acids, glycolaldehyde, and a bile-acid derivative, which show correlation with known tumor promoting host genes. Some features of particular interest due to prior evidence include the correlation of *RBP4* expression (CRC biomarker [42]) with microbiome production of deoxycytidine (DNA building block), *GGT6* with the amino-acid L-Histidine [43], *BMPR1B* with the bile acid derivative 12-Deydrocholate another known cancer promoter [44–47], the blood type gene *ABO* with the amino acid L-Asparate [48], and *PTPN3* with L-Tryptophan, a metabolite implicated in immune-invasion [49]. We also found correlations that suggest beneficial effects of members of the microbiome and their metabolites, such as the production or uptake of resistant starch (a butyrate precursor) and nicotinate (a vitamin B3 precursor), both of which are known to oppose tumorigenesis [50–53]. In addition, oxalate metabolism in the gut microbiome is negatively associated with calcium/ROS channels (*TRPM2*) and epigenetic enzymes (*KDM8*), which is consistent with a protective effect.

Having observed that the expression levels of many host metabolic genes was correlated with the predicted growth rates and metabolic activity of microbes in our models, we wanted to predict whether the changes to a shared growth medium driven by host metabolic activity could directly influence microbial growth patterns. To this end, we performed an *in silico* spent growth medium experiment wherein we constructed models of metabolic reactions for normal tissue and tumor colonocyte cells “grown” on a medium based on the Virtual Metabolic Human (vmh.life) European Union standard diet, which represents a typical Western diet [38]. We calculated the predicted change in growth medium composition due to host model uptake/secretion and used this changed (spent) growth medium as input for the site- and patient- matched MICOM microbiome models’ growth. Overall, we did not predict significant growth rate differences for the entire community on different media (**Supplementary figure 4B**). However, we predicted significant growth rate differences for one taxon (*Blautia*) between the tumor and normal spent media and tumor and normal microbiome combinations (**Figure 3C**). We predicted that *Blautia* from both tumor (Mann-Whitney U *p*=0.027) and normal (Mann-Whitney U *p*=0.0083) microbiome communities grew faster on tumor host cell spent medium as compared to normal host cell spent medium. This is especially interesting because *Blautia* also showed differences in metabolism in the tumor vs normal community in our LASSO regression analysis, and is a known cancer-associated taxon [54,55]

### Some genes share microbiome correlations in patient data and *in vitro* experimental data

Although our analysis reveals correlations between host gene expression and the microbiome, establishing causality requires experimental manipulation. Therefore, we used an *in vitro* system where colon cancer cell lines (HCT116 and HT29) were directly co-cultured with five human gut microbiome communities, allowing us to measure the immediate host transcriptional responses to microbial exposure and identify which associations might represent direct microbial effects on host cells (**Figure 4A**) [26,56,57]. We found significant associations between microbial abundance and the expression of 30 genes involved in host metabolism (q-value<0.1, using generalized linear model controlling for the cancer cell type and other confounders; see **Methods**). These genes were associated with 24 out of 34 genera present, with the most frequent associations involving *Bilophila, Anaerotruncus, Escherichia, Holdemania, and Streptococcus*. Interestingly, several of these genera contain known CRC-associated pathogens, including *E. coli* and *Streptococcus gallolyticus* [58,59] (**Figure 4B)**.

**Figure 4.**
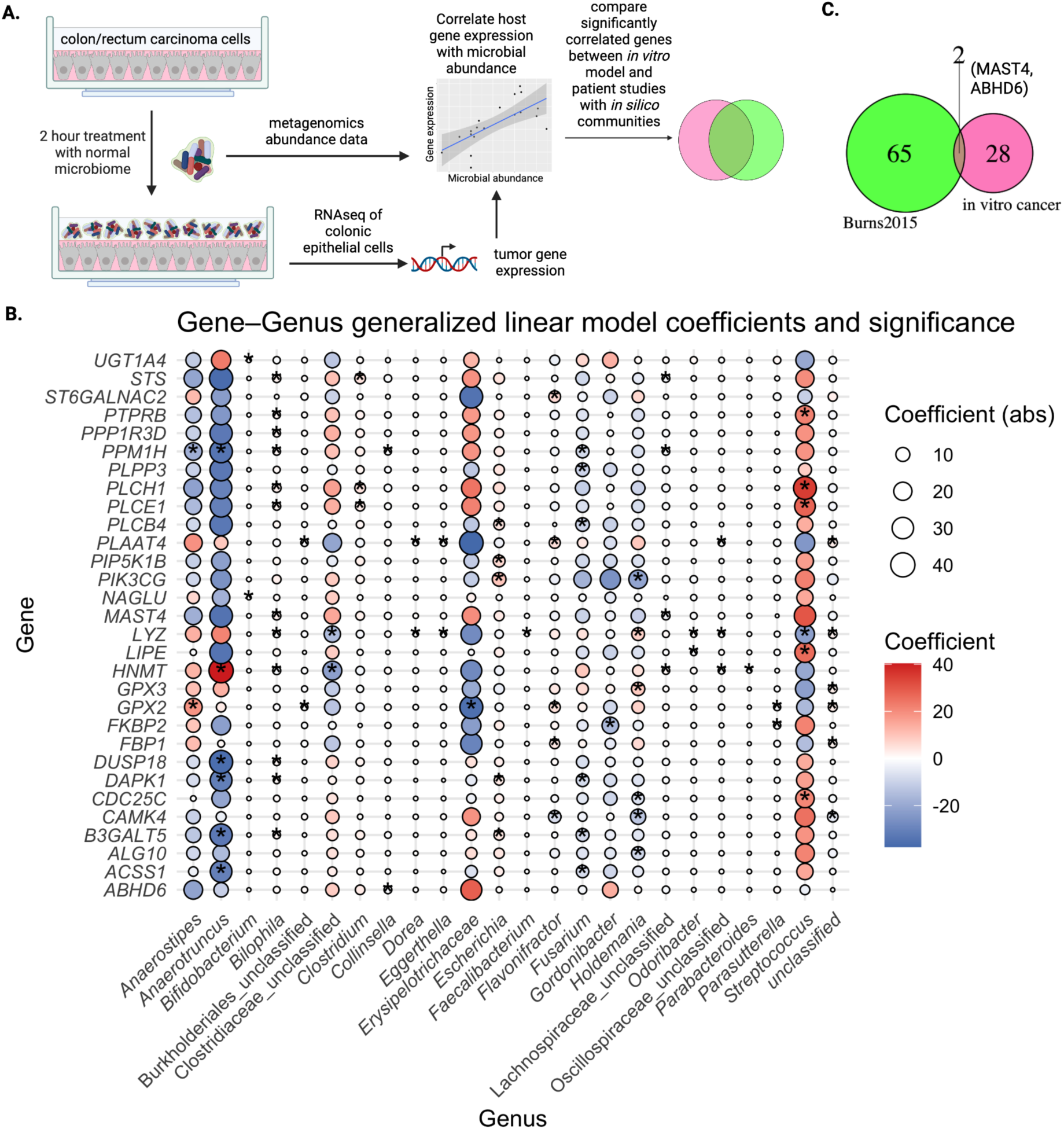
Measuring host-microbiome interactions from *in vitro* microbiome co-culture model. **A.** Schematic of the experimental design for *in vitro* experiments and data collection. Cell lines (HT29 colon carcinoma cells, and HCT116 rectum carcinoma cells) were treated with microbiomes from healthy donors, as described in [27,56]**. B.** Bubble plot of generalized linear model coefficients between host gene expression (columns) and abundance of microbial genera (rows) in the cancer cell lines co-culture experiments outlined in A. **C.** Venn diagrams of shared and unique human metabolic genes showing statistically significant correlations with microbiome features (growth rates in Burns2015 or abundance in coculture experiment) using two tumor cell lines and the *in silico* community models for Burns 2015.

To determine whether our in vitro findings support the associations detected in patient data, we compared the microbiome-associated genes in the two datasets. While the microbial communities differed between the healthy donor samples and CRC patient samples, we could still test whether the same host genes respond to microbiome signals across both contexts. We found that two genes, *MAST4* and *ABHD6*, were associated with the microbiome in both systems (**Figure 4C**). Although the overlap was not statistically significant (Fisher’s exact test, p-value = 0.1769), this likely reflects limited statistical power, as well as the differences between the experimental contexts: the *in vitro* system captures acute responses to healthy microbiomes, while the patient data represents chronic adaptations to tumor-associated communities. Nevertheless, the two genes may reflect relevant microbiome associations. *MAST4* has a positive association with *Bilophila* and unclassified genera in Lachnospiraceae, and encodes a microtubule-associated serine/threonine protein kinase important for cell-signaling. *ABHD6* is negatively associated with *Collinsella*, and has a known protective effect against colorectal cancer through its’ inhibitory effect on the AKT signaling pathway [60].

## Discussion

Overall, our study demonstrates how flux-balance analysis can be used to model the functional characteristics of microbiota across differing growth environments, demonstrating meaningful biological connections between the host and the microbiome. By applying this modeling approach to colorectal cancer patient data, we identified multiple taxonomic and metabolic features of the human microbiome that show links to disease. Using three published datasets of paired tumor and adjacent normal tissue from different CRC patient populations, we built site- and patient- specific metabolic models for the microbial communities present. For one of the studies (Burns et. al 2015), we also used host gene expression to build host metabolic models. We found that *Fusobacterium,* a taxon frequently cited as a correlate and causal factor in CRC carcinogenesis (for review see: [61,62]), is predicted to grow significantly faster in tumor-associated microbiomes as compared to normal tissue-associated microbiomes across all three of the published datasets. In the Burns2015 dataset, we found correlations between modeled microbial growth rates and the expression of host metabolism related genes, as well as associations between the net exchange fluxes for the modeled community and host gene expression. In an *in silico* spent medium experiment most species show no changes in growth rate but *Blautia*, a genus whose growth we also found to correlate with changes in host gene expression, grows preferentially on the metabolic output of CRC host cell models over normal tissue host models. We used an *in-vitro* microbiome-host co-culture model to measure the response of gene expression in different cancer cell lines to the abundance of microbial taxa. While a handful of gene expression changes were shared, this work seemed to capture a different set of genes than the growth-rate based correlations, likely partially due to limited statistical power and the lack of overlap in the composition of the microbial communities tested. Our work represents a novel approach to studying the metabolic interactions between host cells and the microbiome during CRC development and progression, and offers several insights into this process.

Our model prediction that *Fusobacterium* has a consistently higher growth rate in tumor microbiome communities as compared to normal microbiome communities (**Figure 2**) adds to a large body of evidence connecting *Fusobacterium* to the colorectal cancer microbiome. The higher growth rate in tumor compared to normal environments might be driving *Fusobacterium*’s increased abundance in those locations. A previous metabolic modeling study on the CRC microbiome suggested that *Fusobacterium* benefits most strongly from the specific metabolite profile of CRC tissues, although whether that metabolite profile is driven by the host cells or the microbiome remains unclear [63]. Interestingly, we did not predict a difference in *Fusobacterium* growth rates when the microbial community was grown *in silico* with spent medium from host models (**Supplementary figure 4C**). This suggests that the predicted difference in *Fusobacterium* growth is not driven by pro-carcinogenic changes in host cell metabolism, but by some aspect of the microbial community as a whole. Our model also predicts significantly increased valine uptake in *Fusobacterium* in tumor compared to normal tissue associated communities (**Supplementary figure 5D-F.**) This may shed light on a previous finding that colorectal cancer patients with higher consumption of branched chain amino acids including leucine, isoleucine, and valine experienced an increased mortality risk; while the direct causal link is unknown, the authors hypothesized that this might be due to tumors’ preferential use of branched chain amino acids as energy sources for colorectal cancer cancer cells [64]. Our results suggest another possibility: increased valine consumption by the human host could be feeding *Fusobacterium*, a known contributor to CRC progression. The heightened valine consumption predicted for *Fusobacterium* in tumor samples might be driven by competition with host cells for glucose; *Fusobacterium* use of amino acids as a nutrient source may allow it to grow in the tumor environment, where host consumption of glucose is high [65].

Our integrated analysis of microbial metabolism and host gene expression revealed bidirectional metabolic interactions that may drive CRC progression. Using LASSO regression, we identified associations between predicted microbial growth rates and host metabolic gene expression that align with known CRC biology (**Figure 3**). Notably, increased growth of Porphyromonadaceae correlated with upregulation of inflammatory and epigenetic regulatory genes including *MTMR3*, *DNMT3A*, and the tumor-promoting *CBR1*. The Porphyromonadaceae family is consistently overrepresented in colorectal cancer patients, especially the genus *Porphyromonas* and the oral pathogen *Porphyromona gingivalis* within it [41,66,67]. This suggests that certain microbes may thrive by exploiting the inflammatory tumor microenvironment while simultaneously reinforcing it. Supporting this hypothesis, our in silico spent medium experiments suggested that *Blautia*, another CRC-associated genus, preferentially grew on metabolites produced by tumor versus normal host cells (**Figure 3B)**, indicating direct metabolic feeding of pathobionts by cancer cell metabolism [54,55].

The metabolite flux analysis revealed a duality in microbiome-host metabolic exchange. Pro-tumorigenic signatures included correlations between established oncogenes (*AMACR*, *PLOD2*, *ENPP1*) and microbial production of amino acids and simple sugars, consistent with the heightened nucleotide demands and glycolytic stress characteristic of tumor metabolism [68] [69] [70]. The correlation between bile acid derivatives and host gene expression aligns with their known role in CRC development through microbiome-mediated metabolism. Conversely, we identified a protective metabolic counter-signature: microbial metabolism of resistant starch (a butyrate precursor) and nicotinate (vitamin B3 precursor) correlated with downregulation of tumor-promoting genes including *SRC* and *VRK2* [71–75]. This duality suggests that while some microbial metabolic activities support tumorigenesis, others may represent untapped therapeutic opportunities through dietary or probiotic interventions that enhance protective metabolite production. Together, these findings paint a picture of the tumor microenvironment as a metabolically dynamic ecosystem where host and microbial metabolism are closely linked, with implications for both disease progression and potential intervention strategies.

While existing databases are the current gold standard for building microbial communities, there may be strain-specific metabolic features in the real microbiome communities of our patients that are absent from our models, or present in our models but absent in the real community. To ensure as many metabolic reactions as possible were present in our models, we created OTU-specific metabolic models by combining multiple species-level models for the same genus; in this way, we aimed to capture the widest possible metabolic potential of that species. However, rare or understudied metabolic reactions may not be present in published models. Additionally, because we used previously published datasets, we were limited by the lack of multi-omics data. Future studies should experimentally test and validate predictions about growth rates and fluxes of the taxa present in these communities, as well as predictions about the metabolites present in the tumor microenvironment. Since patient gene expression data was only available for Burns2015 [31], we were also unable to validate our host gene expression-microbiome correlations in other patient datasets. Nevertheless, we used an *in vitro* approach to compare patient data to host gene expression changes induced by the microbiome confirming the impact of the microbiome on many of the genes we observed to respond *in silico*. Finally, our study represents a single snapshot of patient data, and our metabolic modeling makes the assumption that the community is a static, steady-state system. While this work provides important insights on the role of specific metabolite exchanges in the microbiome and associations between host gene expression and the microbiome, cancer is a rapidly evolving and changing disease, making temporal dynamics an important part of investigating its causes and progression. We hope that future studies can address this limitation through incorporating longitudinal data, which can shed light on the metabolic activity of the microbiome and host throughout colorectal cancer progression.

In conclusion, our study is the first integration of microbiome data with human gene expression through metabolic modeling in colorectal cancer, a disease known to cause changes in human cell metabolism and suspected to have important metabolic effects in the microbial community. We found evidence supporting many previously suspected associations of the growth of particular taxa with colorectal cancer and human metabolic gene expression, and we also identified some new associations that represent intriguing targets for further research. The approach can be adjusted to other diseases and datasets with many exciting opportunities for future study.

## Acknowledgements

We thank Jeremy Chacón for assistance with setting up the metabolic modeling. We also thank Laura Grieneisen, Kelsey Johnson, Rich Abdill, Mark Swanson, Mattea Allert, Trevor Gould, Samantha Graham, Sabrina Arif, Ashwin Chetty, JJ Colgan, Elizabeth Gibbons, and Karen Tang for their insights and discussion throughout this project.

This work was supported by NIH grant R35GM128716 to R.B, and by a Minnesota Colorectal Cancer Research Foundation to E.M.A. We thank MCCRF for their dedication to supporting research on colorectal cancer.

E.A. and R.B. conceived this project and designed the study. F.L. designed and supervised the in vitro experiments and their data analysis. A.M and A.A designed and conducted the *in vitro* experiments. E.A, S.P., A.M., and K.D.L analyzed and interpreted the data. E.A., R.B., and K.D.L drafted the manuscript. All authors reviewed and approved the final manuscript.

## Methods

### Study design, samples, and data

All patient data used in this manuscript was obtained from previously published studies. Detailed patient information, inclusion criteria, sample collection methods, sequencing protocols, and study information can be found in the original studies. We briefly describe these studies below; further details can be found in **Supplementary table 1**.

In Burns2015 [31], colonic epithelial biopsies were obtained from 44 patients with colorectal cancer. For each patient, samples were obtained both from tumor tissue, and from adjacent normal tissue. 16s rRNA sequencing and host RNAseq were performed for each site, resulting in four sample datasets per patient: normal and tumor-associated 16s rRNA sequences for microbiomes, and normal and tumor tissue host gene expression. We downloaded 16s rRNA OTU read count tables from the SRA (BioProject PRJNA284355). Samples for the original study, which were used to generate these microbiome and host transcriptome data, were obtained through the University of Minnesota Biological Materials Procurement Network (Bionet), and the study was approved by the University of Minnesota Institutional Review Board, protocol 1310E44403. We also obtained the associated RNAseq data from the SRA (BioProject PRJNA816986) [25].

For Hale et al. 2018 [32], 16s sequencing data was obtained from the SRA (BioProject accession PRJNA445346). This study was approved by the Mayo Clinic Institutional Review Board (IRB# 14-007237 and IRB# 622-00). We limited our analysis to a subset of all patients included in this study, according to the following criteria: 1) Only rectum/colon biopsy samples were included (no appendix biopsies), and only samples where both tumor and normal tissue from the same patient were obtained. 2) If tumor and normal samples were obtained from the same patient but different sites in the intestine, that patient’s sample was excluded. 3) If multiple tumor and normal samples from different tissues were obtained from the same patient, we only included colon samples. This resulted in 92 samples from 46 patients; detailed patient information can be found in **Supplementary table 1**.

For Niccolai et al. 2020 [33], colonic biopsies were obtained from 45 patients undergoing surgery at the Careggi University Hospital in Italy in 2018; however, microbiome sequence data for paired normal and tumor tissue samples were only obtained for 40 patients, which is the subset we included in our study. Detailed patient information can be found in **Supplementary table 1**. The 16s rRNA count tables were downloaded from the NCBI Gene Expression Omnibus (GSE163366).

### Preparation and analysis of OTU tables

For all studies described above, we downloaded raw OTU/ASV read count tables as well as associated patient metadata (**Supplementary table 1**). We then removed known contaminant taxa (see **Supplementary table 2** for these) and calculated relative abundances. We filtered out taxa which did not reach a 0.1% threshold in at least one sample. Removing contaminants and abundance filtering did not show any significant differences between tumor and normal communities. We then used the phyloseq package in R [76] (version 1.40.0) to create a phyloseq object from our OTU tables using the *phyloseq()* function. We used the microbiome package (version1.18.0) [77] to calculate Aitchison’s distance beta diversity on our datasets [78]. We chose Aitchison’s distance as a beta diversity metric as it better accounts for the compositionality of microbiome data than other distance metrics [79]. First, we performed a centered-log ratio (CLR) transform of our data using the *microbiome::transform()* function, then used the phyloseq *phyloseq::ordinate()* function to calculate redundancy analysis (RDA) distance matrices on the transformed data. We then performed a permutational multivariate analysis of variance (PERMANOVA) on the distance matrix to examine whether tumor or normal site had an impact on microbiome composition. The PERMANOVA was performed using the *adonis* function in the R package vegan [80] v2.6.2, with 999 permutations. **Figures 1B-D** show the results of this analysis.

### Building metabolic models of microbiome communities using MICOM

Next, we used the taxonomy from these samples to assign each OTU/ASV to a genome-scale metabolic model. We downloaded all bacterial genome-scale metabolic models from the AGORA and MAMBO databases, which are reconstructions based on whole genome sequences and experimental evidence [36,37] and assembled taxonomy information for each database. First, we removed all OTUs/ASVs which were not identified to at least the family level (i.e. order or above). Then, we searched for models in the AGORA database which matched the full taxonomic identification (Domain-Family or Domain-Genus, as available) of the OTUs/ASVs in our study. Since models are built from whole genome sequences, they are identified to the species or strain level. As such, there were often multiple models assigned to each taxon. In these cases, we retained all models and merged them into a single model representing that OTU/ASV using the *micom.util.join_models* function of MICOM [34]. This function takes as input a list of COBRA metabolic models and merges them into a single model by taking the first model in the list, and iteratively adding reactions from subsequent models which are not present in the first model. The result is a COBRA model that contains all metabolic reactions present in each of the input models. We chose this option, rather than selecting a representative model for each OTU/ASV, as it would keep the metabolic potential for that taxon as broad as possible and ensure we were not biasing our results based on an arbitrary selection of a representative model for a given genus.

Briefly, each metabolic reaction present in a cell (as identified by a genome sequence encoding the enzyme that carries out that reaction) is represented as a numerical matrix of stoichiometric coefficients for that reaction. Metabolites that are consumed have a negative coefficient, and metabolites that are produced have a positive coefficient. Each row of the matrix represents a unique metabolite, and each column represents a unique reaction. The matrix is then used to solve for an objective function, which is usually biomass, using linear programming algorithms such as COBRA [28].

For taxa that did not match a model in the AGORA database, we searched for models in the MAMBO database which fully matched the taxonomy (Domain-Family or Genus, as available) of the OTUs/ASVs in our study, again keeping all models when multiple models matched. For the remaining taxa lacking a model, we then searched both AGORA and MAMBO databases for single-taxonomic level matches (i.e. matching ONLY at the family or genus level). We did this to account for the fact that most OTU/ASV tables are built using a specific version of a 16s rRNA database, and taxonomies change across different database versions. For example, the *Collinsella* class was identified as Coriobacteriia in the AGORA model taxonomy and Actinobacteria in Niccolai et al. 2020; all other taxonomic level identifiers were identical. For any taxa in our ASV/OTU tables which still lacked a model, we used CarveMe [81] to generate genome-scale metabolic models from published annotated genomes. These models are described in **Supplementary table 3** and included in the **supplemental data files**. To do this, we searched the NCBI Genome database for annotated whole-genome sequences from a given taxon’s genus. Upon finding the sequence, we downloaded it and used it as input for CarveMe to generate a genome-specific model. If multiple matching genomes were found, we downloaded all genomes obtained from samples of human body sites and used CarveMe to generate models. We then merged the models using MICOM as described above. If no genome from that genus could be obtained, we searched for genomes matching the taxon’s family-level taxonomic identification. If those could not be found, we removed that taxon from our analysis. Detailed information on how many taxa were removed/assigned at each of these steps is found in **Supplementary table 4**. After models had been assigned to all OTUs/ASVs, we used the MICOM Community function to build relative abundance-scaled metabolic models for each community. These communities were saved as .pickle files and can be found in the **supplemental data files**.

Next, we performed flux balance analysis on our communities, using a standardized set of metabolites as *in silico* growth medium. To generate this medium, we first calculated the minimal medium required for each community using MICOM’s *minimal_medium* function. This calculates the smallest set of import fluxes that allow for optimal community growth. To this minimal medium, we then added all components of a European Union standard diet (downloaded from vmh.life, [38]) that could be imported by the taxa in our community models. If a medium component was present at different levels in the community model minimal medium and the EUstandard diet, we retained the higher value. We then assigned the resulting growth medium to the community model and ran the MICOM *cooperative_tradeoff* function with fluxes set to True, and parsimonious FBA set to False. The *cooperative_tradeoff* function in MICOM allows the user to balance maximum community growth (which favors the non-biologically-feasible high growth of a few taxa) with the growth of the maximum number of taxa (which causes lower overall community growth). With one of our datasets [31], we tested several different tradeoff parameters to find one which balanced these two factors (**Supplementary Figure 1**). We found that changing the cooperative tradeoff parameter did not impact differences in community growth rate between tumor and normal tissue samples, but did impact the number of taxa in each community which were able to grow. Since we do not have quantitative growth rates for any of the taxa in our studies, we compared *in silico* growth rates obtained in MICOM to previously published quantitative growth rates from similar studies [82]. We defined bacterial growth as at least 10^-6^/h. Based on these results, we found that a cooperative tradeoff value of 0.2-0.25 was optimal for the communities in our study. If we chose a higher tradeoff value, fewer than a third of the taxa in our communities would grow, and their growth rates would be far higher than is biologically feasible. If we chose a tradeoff value below 0.2, however, the *cooperative_tradeoff* function would fail to find a solution, indicating model infeasibility.

### Analysis of MICOM outputs

All analyses were performed in Python version 3.7.6 or RStudio version 2021.09.2 as specified. Statistical analyses were performed in RStudio.

From the *cooperative_tradeoff* function in MICOM, we obtained several outputs for each patient-and site-specific *in silico* microbiome community. These included community and individual species growth rates for each community; community-level measures of metabolite uptake and excretion flux; and taxon-specific measurements of flux for all reactions included in taxon-specific metabolic models. We used RStudio version 2021.09.2 to analyze differences in these output parameters between tumor-associated and normal tissue-associated *in silico* microbiome communities. Since the data in our studies are paired, meaning we have both tumor-associated microbiome samples and normal tissue-associated samples, all differences calculated were within-patient differences and all statistical analyses performed were for paired data.

### Differential growth and metabolic flux in three datasets

To analyze differences in individual species’ growth rates between tumor and normal samples, we combined the results from all community runs and performed several filtering steps to focus our analyses only on results with a metabolic implication in the host. If a taxon’s growth rate from the model was less than 10^-6^, we defined that taxon as non-growing, as recommended in the MICOM documentation [34]. We then collapsed the growth rates across genera, wherein we assigned the growth rate to a given genus within a sample to the average growth rate of all OTUs/ASVs from that genus. Upon calculating the difference in growth rate of each taxon between tumor and normal sites within a patient, we removed taxa where the difference in growth rates between tumor and normal samples was zero (i.e. growth rates were identical). We reasoned that taxa with the same growth rates were unlikely to contribute to CRC-related metabolism. We then removed taxa which did not appear in at least 50% of all samples, to ensure we were capturing microbial taxa which were present and active in the majority of patients within a study. **Supplementary table 5** outlines the number of taxa left after each step of this filtering approach. For each genus, we then performed a Wilcoxon signed-rank test with False Discovery Rate-corrected *p*-values to examine whether growth rates differed significantly between tumor and normal tissue associated communities across patients.

To analyze differences in metabolic fluxes between tumor and normal samples, we performed similar filtering steps. First, we removed all flux pathways that did not reach a threshold of at least 10^-6^ in at least 10% of OTUs/ASVs in a given dataset. We also only analyzed exchange fluxes (i.e. fluxpaths which involved exchanging a metabolite between the interior of the *in silico* model cell and the exterior media environment), as the activity of these pathways is most likely to impact the host cells, which share a growth medium environment with the microbiome. Since we were most interested in the total activity of a given fluxpath in the community, rather than the activity of individual taxa, we then summed the activity of each exchange fluxpath across taxa within a given sample community. This gave us a net fluxpath activity value for each fluxpath present in the community. We then filtered out fluxpaths which had identical activity levels between tumor and normal samples within a patient, and those which were not present in every sample in a given study. **Supplementary table 6** outlines the number of fluxpaths left after each step of this filtering approach. For each fluxpath, we then performed a Wilcoxon signed-rank test with False Discovery Rate-corrected *p*-values to examine whether fluxpath activity differed significantly between tumor and normal tissue associated communities across patients. The results for this analysis can be found in **Figure 2 B-F**.

### Building host models using CORDA

To build patient-specific models of host tissue (applicable for Burns 2015 dataset only, as this was the only patient dataset that included gene expression data from both host and microbiome samples), we used a program called CORDA (Cost Optimization Reaction Dependency Assessment) [35]. Briefly, this program uses gene expression or protein production information to “prune” a generalized human metabolic model, thereby creating a patient- and tissue-specific metabolic model of a human cell. To create these models, CORDA performs a dependency assessment of all reactions present in a generalized human model, for which we used RECON3D [83]. The dependency assessment is based on reactions which have been sorted into high/medium/low/negative confidence groups, defined by the user from gene expression data. When the patient-specific model is built, CORDA includes all high confidence reactions and as many medium confidence reactions and as few low confidence reactions as possible.

The first step in using CORDA to create patient- and site-specific host cell models for our study was to re-analyze Burns 2015 RNAseq data and use Boolean mapping to sort gene expression into five categories: high/medium/low/none/not present. First, we downloaded the data from the SRA (BioProject PRJNA816986) [25] and calculated abundance estimates (transcripts per million, TPM) using kallisto [84]. We then binned the expression data based on the criteria suggested by Expression Atlas. Genes with TPM <=0.5 were assigned −1 (no expression); genes with TPM 0.5-10 were assigned a score of 1 (low expression); genes with TPM 11-1000 were assigned a score of 2 (medium expression), and genes with TPM> 1000 were assigned a score of 3 (high expression). Genes not included in the dataset were assigned a score of 0. The number of genes included in each expression bin per patient is found in **Supplementary table S7**.

Next, we obtained all gene-protein-reaction associations (GPRs) from the RECON3D generalized human metabolic model. GPRs form the basis of the metabolic model, and describe the connection between gene expression and metabolic pathway activity. By default, these GPRs take the form of and/or Boolean rules, such that the reaction is considered to occur if certain genes are expressed. For example, a reaction *a* to produce compound A might require three enzymes, Q, R, and S. The genes encoding Q, R, and S must all be expressed in order for compound A to be produced. The GPR for reaction *a* is therefore Q AND R AND S. To use CORDA, all AND Boolean associations from RECON3D were replaced by MIN, and OR Boolean associations were replaced by MAX. The expression bins corresponding to all genes were then substituted into the GPRs, resulting in Boolean rules of −1/0/1/2/3 for each GPR in the model. These rules corresponded to five different confidence levels for these reactions in the model: not present/excluded from model (−1); unknown confidence, may be included if necessary (0), low confidence, may be included if necessary (1), medium confidence, should be included if necessary (2), and high confidence, must be included if possible (3). Importantly, like most metabolic models, RECON3D includes several reactions without associated GPRs which are required for the model to run. These include reactions which are thermodynamically required for cellular growth, but the genes responsible for this reaction are unknown [83]. These reactions were assigned a Boolean confidence level of 3, as the model does not run without them. Additionally, five parameters are required for CORDA as described in more detail in the paper. We used the default values for all parameters. We used the *opt_build* function of CORDA to generate patient- and site-specific metabolic models for the Burns 2015 data; these models can be found in **supplemental data files**. From these models we obtained data on growth rates, imported/exported metabolites from the model, and flux of all metabolic pathways. We analyzed these data in R as described below.

### Combining microbiome and host cell models

To investigate how host consumption of growth medium metabolites impacts the microbiome, we performed an *in silico* spent growth medium experiment. *In vitro* spent medium experiments involve growing a primary culture of cells, filtering out cellular components such that only secreted factors and growth medium components are left, and using the filtrate to grow a secondary culture of cells [85]. To recapitulate this *in silico*, we calculated the change in metabolite levels before and after running flux-balance analysis in CORDA on each patient- and site-specific host cell model. Metabolites which were taken up by the cell model were subtracted from the original growth medium based on the standard european diet, and metabolites which were excreted were added to the growth medium. We then used this spent medium as input for the patient- and site-matched microbiome model, and reran MICOM as described above. We also performed the spent medium analysis in reverse (first running MICOM, collecting the spent medium from the *in silico* microbiome and using it as growth medium for the *in silico* human cells).

### Integration of gene expression and metabolic modeling data in Burns 2015

For the Burns 2015 dataset only, we had host gene expression data from both tumor and normal tissue sites, allowing us to integrate them in a LASSO regression model following a previously published in-house pipeline [25].

We downloaded the published raw RNAseq data from NCBI and ran kallisto [84] to quantify transcript abundances. We next filtered for genes that are present above a threshold of 9 reads in at least half the samples and are involved in metabolism. For filtering by metabolism-related genes, we downloaded a list of all metabolic reactions and their associated human genes from HumanCyc [86], and extracted a nonredundant list of genes (see **Supplementary table 7** for this list). This list included 2,477 metabolic reactions, and 2,629 genes, 2,137 of which were present in our dataset at sufficient prevalence and abundance. Next we calculate log2 fold change in expression between normal and tumor samples of these genes using DESeq2 [87].

Similarly to the gene expression data, we prepared the output of MICOM metabolic models by creating a combined table of metabolic flux and growth rate microbial features. For each microbial metabolite exchanged with the medium in at least 22 of the 44 samples, we summarized the fluxpath activity across the entire microbiome as a sum of microbial species-specific fluxes. We then calculated the change in fluxpath activity between the normal tissue model and tumor tissue model for each patient and included it as a microbial feature. Similarly, for the microbial growth rates, we calculated the change in growth rate from normal to tumor samples for features at the family, genus, and species level and filtered for taxa that show change in more than 1/4 of the patients. This resulted in a microbial feature table consisting of 27 changes in growth rate and 138 changes in metabolic fluxpath activity. We then normalized this table using the base R ‘scale’ function. We searched for biologically meaningful associations between changes in microbial features (fluxes or growth rates) and changes in human gene expression using an in-house developed machine learning-based multi-omics integration framework [25] using lasso penalized regression [39].

### The LASSO penalized regression model

We used LASSO penalized regression using the R package glmnet (version 4.1-8) to identify pairwise associations between the change in expression of individual host genes and the change in either *in silico* growth rate of microbial taxa or the change in *in silico* metabolic flux between the microbiome and medium. This LASSO penalized regression is done in a gene-wise manner where the change in gene expression is the response variable and the change in microbial features (growth rates and metabolic flux rates) are predictor variables. We expect only a small number of the many microbial features to correlate with any given gene’s change in expression and the L1 regularization in LASSO regression does the variable selection by automatically setting the LASSO coefficient of irrelevant microbial features to 0. The LASSO regression coefficient β is estimated by minimizing the following equation.

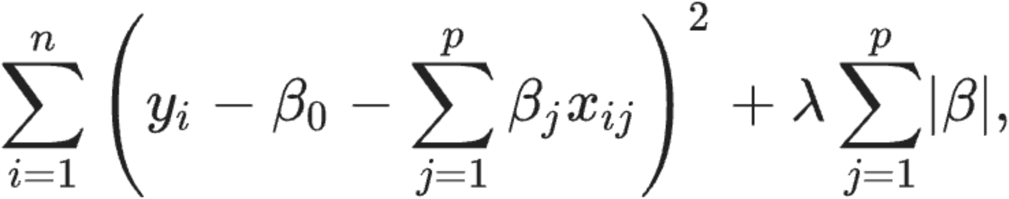

𝑛 is the number of samples (44), where 1 ≤ 𝑖 ≤ 𝑛, 𝑝 is the number of predictors (change in growth rate and metabolic flux), where 1 ≤ 𝑗 ≤ 𝑝, 𝑦 is the response (change in gene expression), 𝑥 is the predictor (change in growth rate or metabolic flux), and 𝜆 is the tuning parameter, where 𝜆 ≥ 0.

The first term in this equation is the residual sum of squares, so the amount of variance in the response variable (change in gene expression) that can not be explained by the predictor (change in microbial features). The second term is the 𝑙_!_norm of the coefficients, which depending on the selected 𝜆, forces variable selection. To estimate 𝜆, we used leave-one-out cross validation.

To obtain the 95% confidence interval and p-values for the regression coefficient 𝛽 of each microbial feature, we used a regularized projection approach known as desparsified LASSO using the ‘hdi’ R package (version 0.1-9) [88].

To make sure the microbial features were robustly associated with the changes in host gene expression we implemented stability selection [89]. Briefly, we selected random subsets of the data, fit the LASSO model to this subset with a penalty close to the estimated best one, and record the microbial features that show association with the gene expression changes. We do this 𝐾 times (in this case 10) and only select the microbial features that are selected a pre-specified fraction of times. The LASSO results can be viewed in **supplementary table 9**.

### Colon cancer cell line preparation for *in vitro* host-microbiome model

The experimental protocol used for the treatment of colon cancer cell lines (HCT116 and HT29) with microbiota has previously been described in Richards et al., 2016 [26]. The cells were cultured on plates on or flasks coated with poly-l-lysine (PLL), according to the supplier’s specifications (ScienCell 0413). Cells were cultured in colonic epithelial cell medium supplemented with colonic epithelial cell growth supplement and penicillin-streptomycin according to the manufacturer’s protocol (ScienCell 2951) at 37°C with 5% CO_2_. 24 hours before treatment, cells were changed to antibiotic-free medium and moved to an incubator at 37°C, 5% CO_2_, and a reduced 5% O_2_.

### Microbiome sample acquisition and preparation for *in vitro* host-microbe model

Human fecal samples were purchased from OpenBiome and arrived frozen on dry ice. The following briefly describes the protocol used by OpenBiome to process stool samples. The samples were collected by OpenBiome and given to a technician within 1 hour of defecation. The mass of the sample is measured and transferred to a sterile biosafety cabinet. The stool sample is put into a sterile filter bag and a sterile filtered dilutant of 12.5% glycerol is added with a normal saline buffer (0.90% [wt/vol] NaCl in water). The sample solution is then introduced to a homogenizer blender for 60s and aliquoted into sterile bottles. The bottles are then immediately frozen at −80°C. Any sample not fully processed within 2 hours of passage is destroyed.

Fecal microbiota were not thawed until the day of the experiment. Prior to treatment, the microbiota were thawed at 30°C and the microbial density (OD_600_) was assessed via a spectrophotometer (Bio-Rad SmartSpec 3000).

### Microbiome-colon cancer cell line co-culture experiment

Medium was removed from the colon cancer cells and fresh antibiotic-free medium was added to the cells, with a final microbial ratio of 10:1 microbe:colonocyte in each well. Additional wells containing only colon cancer cell lines were also cultured in the same 24-well plate for use as controls.

After 2 hours, the wells were scraped on ice, pelleted, and washed with cold phosphate-buffered saline (PBS), then resuspended in lysis buffer (Dynabeads mRNA Direct kit, ThermoFisher Scientific, Waltham, Massachusetts, USA) and stored at −80°C until extraction of host RNA for RNA-seq. We conducted metagenomic shotgun sequencing on the 5 different microbiomes used. Previous experiments have shown that microbiome composition does not change drastically over the 2 hour co-culture period [27] and we thus used only the composition before preparation to run our correlation with gene expression analysis. The experiment for each cell line was performed on separate days.

### RNA-sequencing and data processing for *in vitro* host-microbiome model

Poly-adenylated mRNA was isolated from thawed cell lysates using the Dynabeads mRNA Direct Kit (Ambion) following the manufacturer’s instructions. RNA-seq libraries were prepared using a protocol modified from the NEBNext Ultradirectional (NEB) library preparation protocol to use Barcodes from BIOO Scientific added by ligation as described in [27]. The libraries were then pooled and sequenced on two lanes of the Illumina Next-seq 500 in the Luca/Pique-Regi laboratory at Wayne State University using the high output kits for 75 cycles to obtain paired-end reads. Reads were 80 bp in length. Read counts ranged between 12,632,223 and 36,747,968 reads per sample, with a mean of 18,726,038 and median of 16,993,999 reads per sample.

FastQC was used to determine the quality of reads from the raw data (FASTQC, version 0.11.5). Gene expression data was quantified using the CHURP pipeline (v 0.2.2) [90]. Reads were trimmed using Trimmomatic [91]. Duplicates were removed using Picard. Finally, all reads were aligned to the human genome GRCH38 using HISAT2 [92]. As was done for the Burns2015 data, we next filtered for genes that are present above a threshold of 9 reads in at least half the samples as well as for metabolism-related genes, using our list of all metabolic reactions and their associated human genes from HumanCyc [86](see **Supplementary table 7** for this list). This ensures the comparison continues our focus on metabolism-related genes.

### Metagenomic shotgun sequencing for *in vitro* host microbiome model

Metagenomic shotgun sequencing on prepared microbiota samples was performed at the University of Minnesota Genomics Center (UMGC). DNA samples were quantified using a fluorimetric PicoGreen assay. gDNA samples were converted to Illumina sequencing libraries using Illumina’s NexteraXT DNA Sample Preparation Kit (Cat. # FC-130-1005). 1 ng of gDNA was simultaneously fragmented and tagged with a unique adapter sequence. This “tagmentation” step is mediated by a transposase. The tagmented DNA was simultaneously indexed and amplified with 12 PCR cycles. Final library size distribution was validated using capillary electrophoresis and quantified using fluorimetry (PicoGreen). Truseq libraries were hybridized to a NextSeq. Base call (.bcl) files for each cycle of sequencing were generated by Illumina Real Time Analysis (RTA) software. The base call files were demultiplexed and then converted to index specific fastq files using the MiSeq Reporter software on-instrument.

### Characterizing microbiota for *in vitro* host microbiome model

To identify microbial features from the metagenomic shotgun sequencing data, including taxa and pathway abundances, we used the HUMAnN2 pipeline with Metaphlan2 (HUMAnN2 v0.11.1, Metaphlan2 v0.2.6.0)[93,94]. FastQC v0.11.7 was used to determine the quality of sequencing reads before trimming. Sequencing adapters were trimmed from the raw reads using Trimmomatic (Trimmomatic v0.33)[91]. FastQC v0.11.7 was again used to determine quality of sequencing reads after trimming the sequencing adapters from the reads. Metaphlan2 was used to assign taxonomy at all taxonomic levels to the sequencing reads in each sequencing file, and in particular to get relative abundances of microbial taxa for each sample. For further analysis, we used only the raw pre-coculture metagenomics data as the microbiome does not change over the 2h time period [27]

### Finding gene expression - microbial genus abundance correlations in the *in vitro* host-microbiome model

The list of microbial genera was filtered as follows. First we removed the Bacteroides genus since we spiked a pathogenic strain of *B.fragilis* into the microbiomes to test a separate hypothesis and are controlling for its abundance when testing the rest of the genera. Each microbial genus tested has to be present in at least 3 of the 5 microbiomes used, leaving a total of 35 microbial genera to test. The relative abundances were transformed with a centered log ratio transformation. The following generalized linear model was run for each cell type for each of the 36 microbial genera tested:

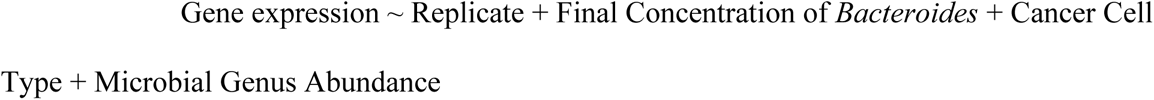

Where replicate indicates the experimental replicate, Final Concentration of *Bacteroides* is a numeric variable indicating the final concentration of *Bacteroides* after spiking in the ETBF strain of *B. fragilis*, Cancer Cell Type the cancer cell line (HT29 colon carcinoma cells or HCT116 rectum carcinoma cells) and Microbial Genus Abundance is the transformed relative abundance of microbial genera present in the microbiome community. Significant host gene-microbial genus correlations were defined at *q*-value <0.1 and reported in **Supplementary table 10**. We used the ggplot2’s geompoint to create a bubble plot of these correlations (**Figure 4B**)

### Comparison between clinical and *in vitro* host-microbe interactions

We used the VennDiagram package in R (version 1.7.3) to generate Venn diagrams showing the overlap in significantly microbial feature-correlated genes between the Burns2015 dataset (changes in host gene expression between tumor and normal sites associated with *in silico* microbial growth rates), and the co-culture dataset (changes in host gene expression associated with microbial abundance cancer cell lines before and after 2 hours of treatment with a healthy microbiome). We used the fisher.test function in the stats R package to test the significance of the overlap.

### Modeling Approach Considerations

For modeling large microbial communities, two main approaches have been used: lumped/”enzyme soup” approaches, wherein the metabolic capabilities of all community members are combined into a single large model; and compartmentalized, wherein separation of microbes and their metabolic capabilities is maintained [95,96]. The “enzyme soup” approach is often necessary when working with metagenomic shotgun sequencing data, where the taxon associated with a given enzyme/reaction may not be known. It is most useful for situations where overall community metabolism, rather than the metabolism of individual taxa, is of interest, and in situations where there is little *a priori* knowledge about the taxa present in the community [97]. However, given that the human colon microbiome is one of the most heavily studied microbiomes [98] and the fact that large databases of curated human gut microbes already exist [36,37], we chose to use a compartmentalized approach. This allowed us to maintain the separate metabolic activities of each taxon in the community, and study taxon-level metabolic and growth activity.

The approach we chose to use, MICOM, combines user-supplied microbial abundance data with database-supplied models to generate static, compartmentalized microbiome models [95]. We chose an approach that relied on previously generated manually curated models for two main reasons. The first was that we wanted to investigate patient-matched tissue biopsy microbiome samples in colorectal cancer, but the low microbial biomass of colonic biopsies makes it difficult to obtain sufficient metagenomic shotgun sequencing coverage to construct *de novo* metabolic models [99]. Second, while algorithms to construct *de novo* metabolic models from MAGs are improving, they remain less accurate than manually curated models incorporating detailed biochemical and growth information [95]. We therefore chose an approach that was broadly applicable across studies, and mostly relied on curated models rather than *de novo* models.

