## Supplementary Figures for "Metabolic modeling and functional genomics reveal taxa and host gene interactions in colorectal cancer"

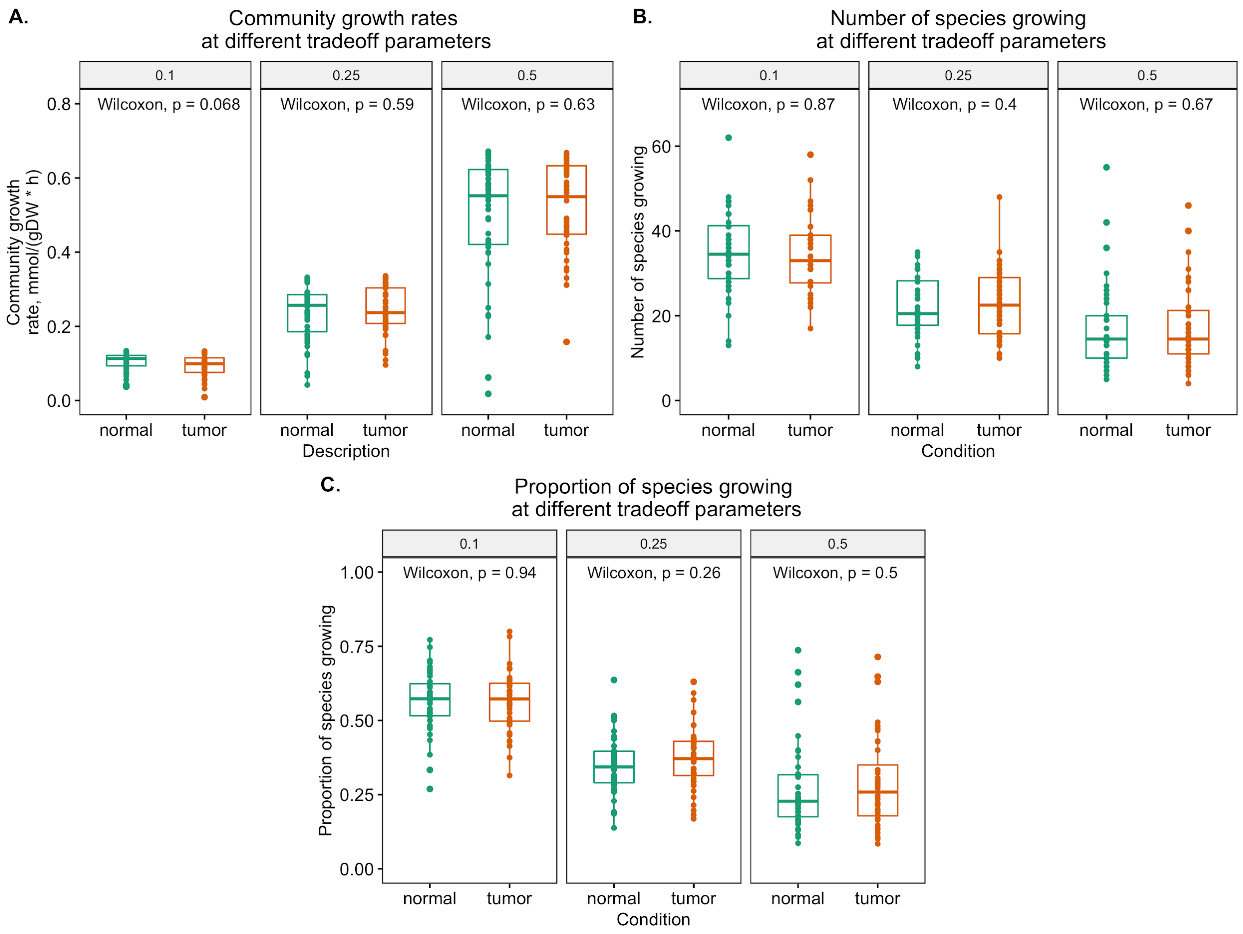


**Supplementary figure 1**. Testing the impact of varying MICOM community growth tradeoff parameters on community growth rates (**A.**), number of species growing per community **(B.**), and proportion of species from each community growing (**C.**) Differences between tumor and normal communities were tested with paired Wilcoxon rank-sum tests.

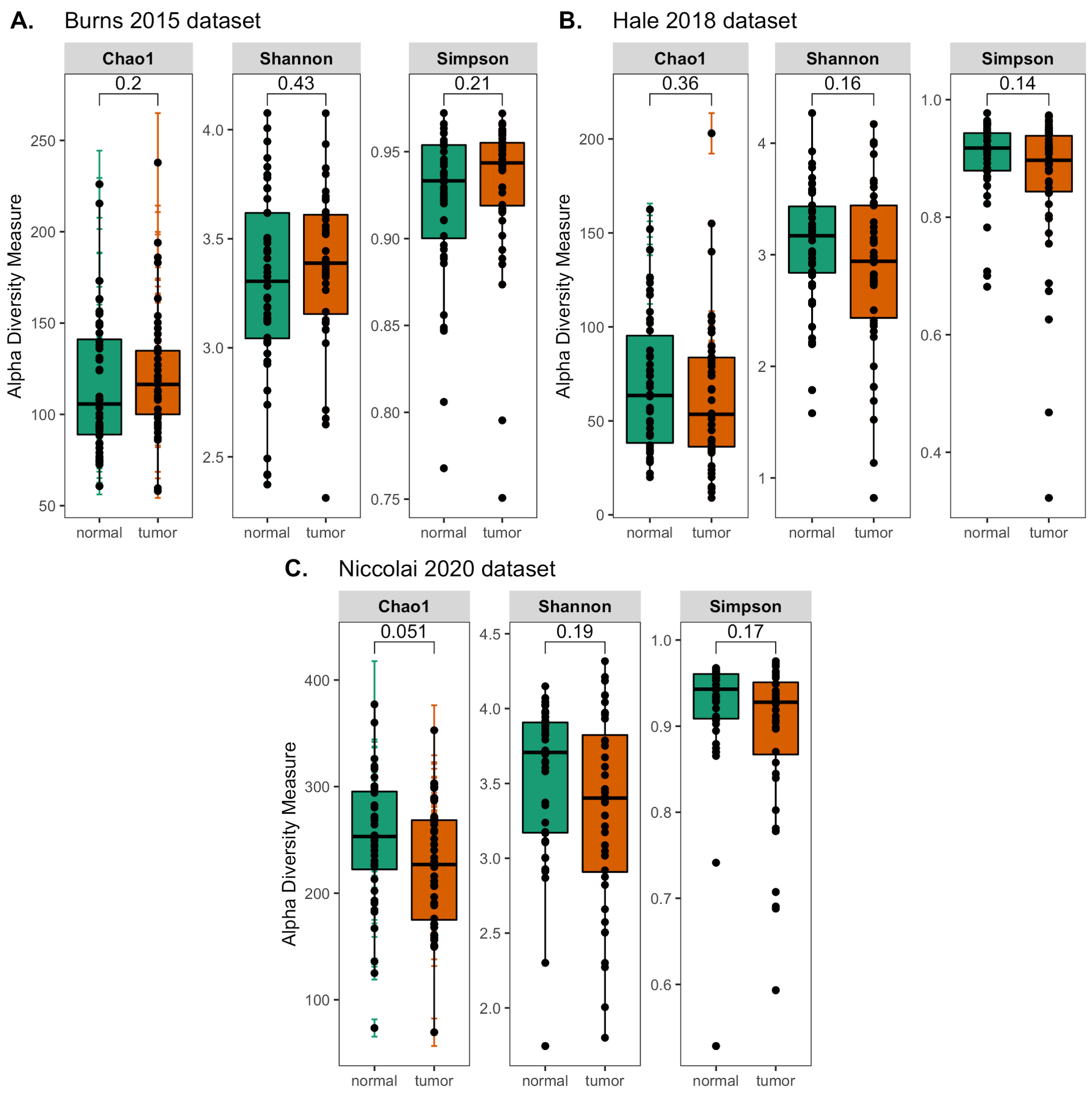


**Supplementary figure 2.** Alpha diversity metrics for OTU tables used in building *in silico* microbiome communities from Burns2015 (**A.**), Hale2018 (**B.**), and Niccolai2020 (**C.**) p-values comparing Chao1, Shannon, and Simpson alpha diversity between tumor and normal samples obtained from paired Wilcoxon rank-sum tests.

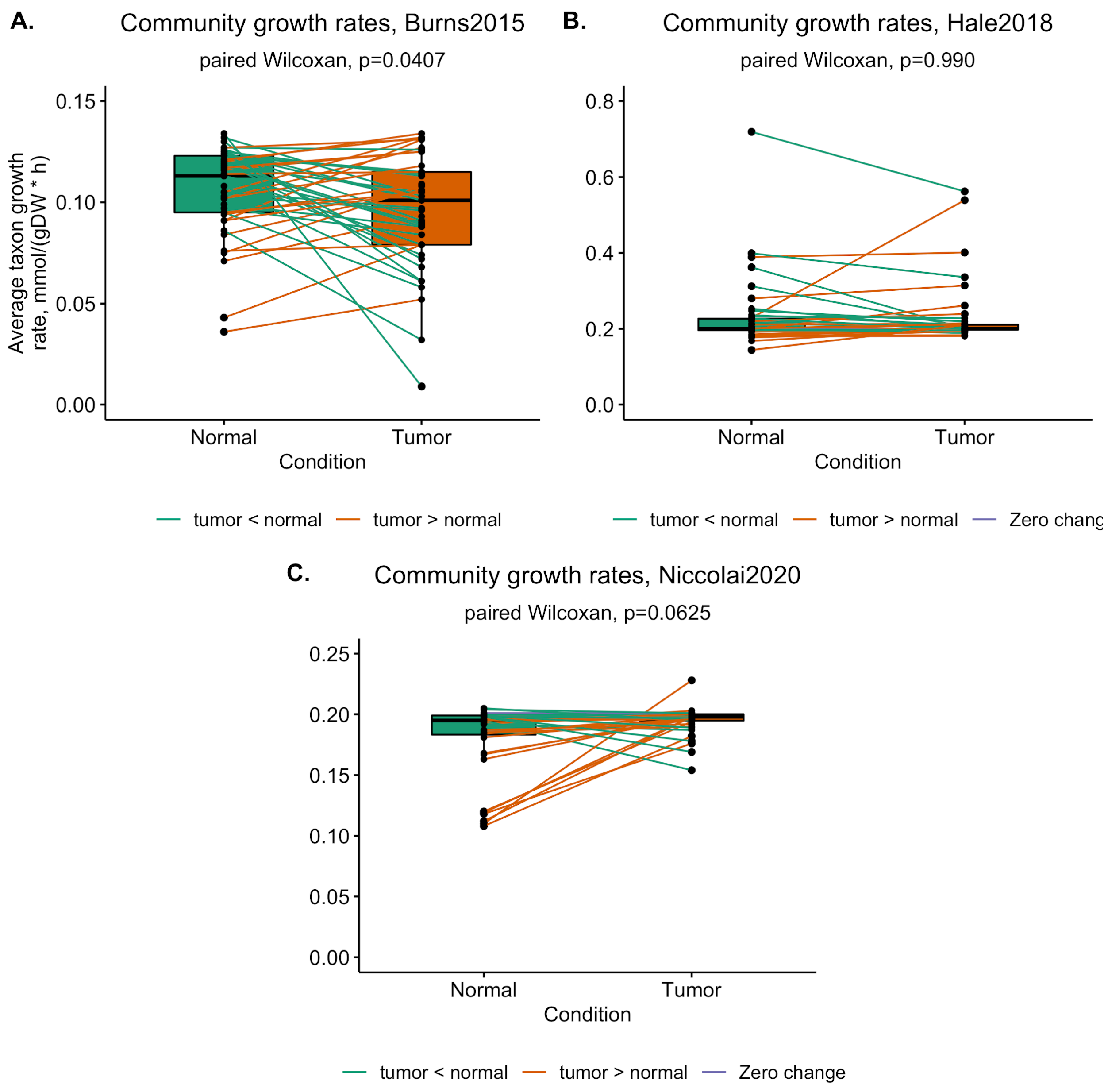


**Supplementary figure 3.** Overall community growth rates in normal tissue vs tumor tissue associated *in silico* microbial communities for Burns2015 (**A.**), Hale2018 (**B.**), and Niccolai2020 (**C.**) datasets. Green lines connect samples from tumor and normal sites in the same patient where the community growth rate was higher in normal tissue than tumor tissue community; orange lines connect samples in patients where the community growth rate was higher in tumor tissue. *P*-values from paired Wilcoxon rank-sum tests.

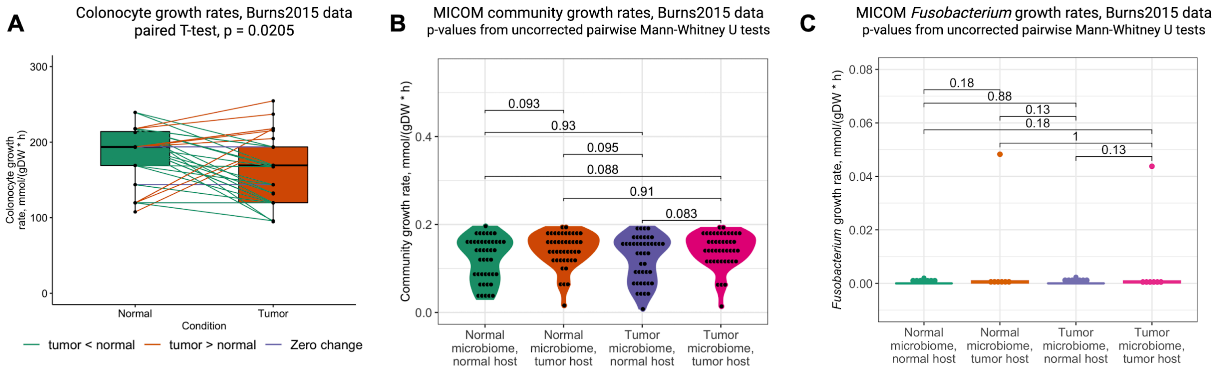


**Supplementary figure 4:** Flux-balance analysis with human cell metabolic models. **A.** CORDA generated growth rates of colonocyte models of normal tissue versus tumor tissue cells. Lines connect tumor and normal tissue models built from RNAseq data from different sites within a patient. Orange lines represent tumor cells growing faster than normal cells, and green lines represent normal cells growing faster than tumor cells. *P*-value from paired Student's T-test. **B.** Community growth rates for normal and tumor tissue-associated microbiomes growing on spent growth medium from normal or tumor tissue host cells. Each point represents the growth rate of a single site- and patient-specific *in silico* microbiome community. *P*-values from Mann-Whitney U tests. **C.** Growth rate of *Fusobacterium* genus in normal or tumor tissue-associated communities grown on spent growth medium from tumor or normal host tissue models. Each point represents the average growth rate of any taxa in the *Fusobacterium* genus growing in a patient and site-specific *in silico* microbiome community. *P*-values from Mann-Whitney U tests.

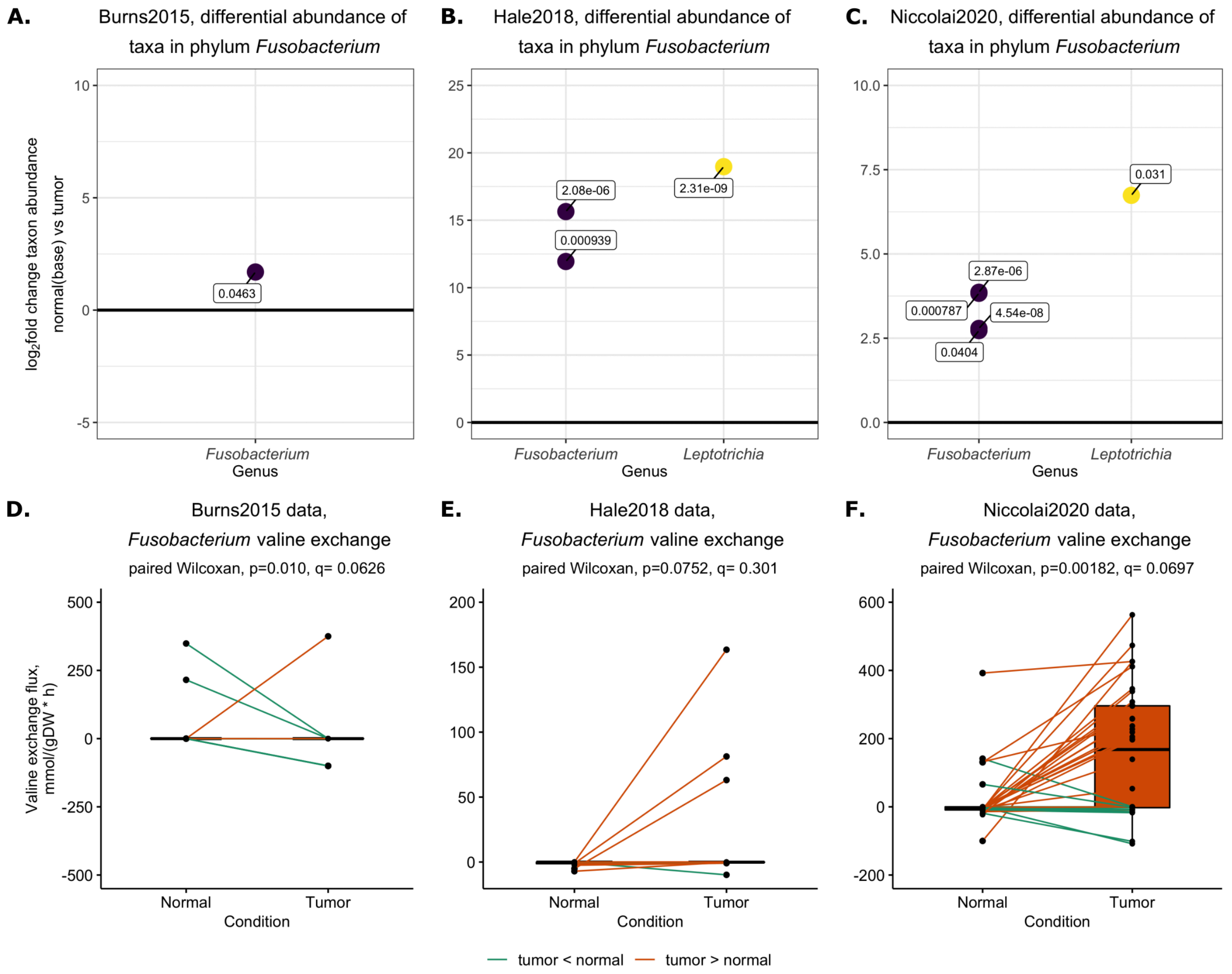


**Supplementary figure 5**. *Fusobacterium* abundance and valine flux across three studies. The relative abundance of *Fusobacterium* in OTU tables from Burns2015 (**A.**), Hale2018 (**B.**), and Niccolai2020 (**C.**). The differential abundance of all taxa within the phylum *Fusobacterium* was identified with DESeq2 and plotted as the $\log_2$fold change of taxon relative abundance in tumor (baseline) versus normal tissue. Points above the dark line represent genera with higher relative abundance in tumor tissue as compared to normal tissue. *P*-values from paired Wilcoxon signed-rank tests with False Discovery Rate correction. **D.-F.** Valine exchange in *Fusobacterium* taxa growing in Burns2015, Hale2018, and Niccolai2020 *in silico* communities, respectively. Positive y-axis (flux) values represent valine import into *Fusobacterium*, and negative values represent valine export from *Fusobacterium* into the growth medium environment. Lines connect *Fusobacterium* growing in the same patient but at different sites (tumor or normal tissue associated communities). Orange lines represent higher import in *Fusobacterium* from tumor communities, and green lines represent higher import in *Fusobacterium* from normal tissue-associated communities. *P*-values from paired Wilcoxon signed-rank tests; *q* values from FDR correction.
